# An Improved Fertilization Protocol for *Ascidiella aspersa*: Promoting a New Chordate Model in Developmental Biology

**DOI:** 10.1101/2025.03.10.642315

**Authors:** Yuki S. Kogure, Kohji Hotta

**Author notes:** Correspondence: Kohji Hotta.

## Abstract

The invasive ascidian *Ascidiella aspersa* is known to be globally distributed (Muller, 1776) and characterized by its ultra-transparent embryos (Shito et al., 2021), which are ideal for studies of developmental biology and bioimaging. However, its use as a model organism has been limited due to the exceptionally low fertilization rate of dechorionated eggs. In this study, we present a new fertilization protocol that drastically increases the fertilization rate from below 10% to 80%, allowing for synchronously developing embryos without polyspermy. This protocol promotes the adoption of ascidian *Ascidiella aspersa* as a new chordate model in developmental biology.

**Summary statement:** Establishment of fertilization protocols for the transparent ascidian *Ascidiella aspersa* as a new model for chordate development studies.

## 1. Introduction

Ascidians are the closest relatives of vertebrates, and their simple body plan and rapid development make them useful model organisms for elucidating the mechanism of genetic events (Lemaire, Smith, and Nishida 2008), morphogenesis (Kogure et al. 2022; Meinertzhagen and Okamura 2001), neuronal circuits (Ryan, Lu, and Meinertzhagen 2017) in chordate development.

In Europe, *Phallusia mammillata*, which has very transparent embryos, has been used as a standard model organism among ascidians. Furthermore, in this species, externally introduced mRNA can be translated early from the egg, making it suitable for imaging approaches (McDougall, Lee, and Dumollard 2014). However, since *Phallusia mamillata* does not inhabit the United States or Japan, there are limits to its widespread use worldwide.

One species of ascidian, *Ascidiella aspersa*, is a globally distributed invasive species (Kanamori et al. 2017) with high transparency from egg to larva (Shito et al., 2020) and early stage translation of external mRNA since the unfertilized egg (Funakoshi et al., 2021; Yasuo and McDougall, 2018). These characteristics make *A. aspersa* an ideal candidate for bioimaging studies during early development, presenting it as a potentially valuable model organism in developmental biology. However, *A. aspersa* faces challenges with fertilization in dechorionated eggs. Dechorionation is essential for various genetic manipulations, such as electroporation and injection. A low fertilization rate in dechorionated eggs has been thought because of contamination from highly acidic body fluids (Bell et al. 1982), as well as polyspermy. Due to these difficulties, *A. aspersa* has not been widely adopted as a model organism for a long time. In this study, we addressed these issues by reconsidering the steps in the fertilization protocol. This improvement represents a breakthrough in establishing *A. aspersa* as a globally recognized model organism.

## 2. Results

### 2.1 Removal of tunic before Isolation of eggs and sperm

One possible reason for the low fertilization rate is thought to be contamination from highly acidic body fluids (Bolton and Havenhand 1996).

Therefore, we took care to remove body fluids at every step.

1. **Removal of the Tunic**: To avoid damaging the gonoduct, begun the incision from the atrial siphon side (indicated by the arrowhead in Fig. 1A) and proceed carefully toward the bottom of the body (Fig. 1B). Then, remove the entire tunic from the body by hand. The gonoduct can be seen as the white line (arrow in Fig. 1C).
2. **Drain of Body Fluid**: The lower part of the body containing the heart region was cut in advance to drain most of the acidic body fluid (Fig. 1D). Furthermore, the remaining surface body fluids were wiped off with paper towels (Fig. 1E) and *A. aspersa* body was placed between paper towels for a few minutes to absorb fluids (Fig. 1F) minimizing body fluid contamination when collecting the eggs.

**Fig. 1.**
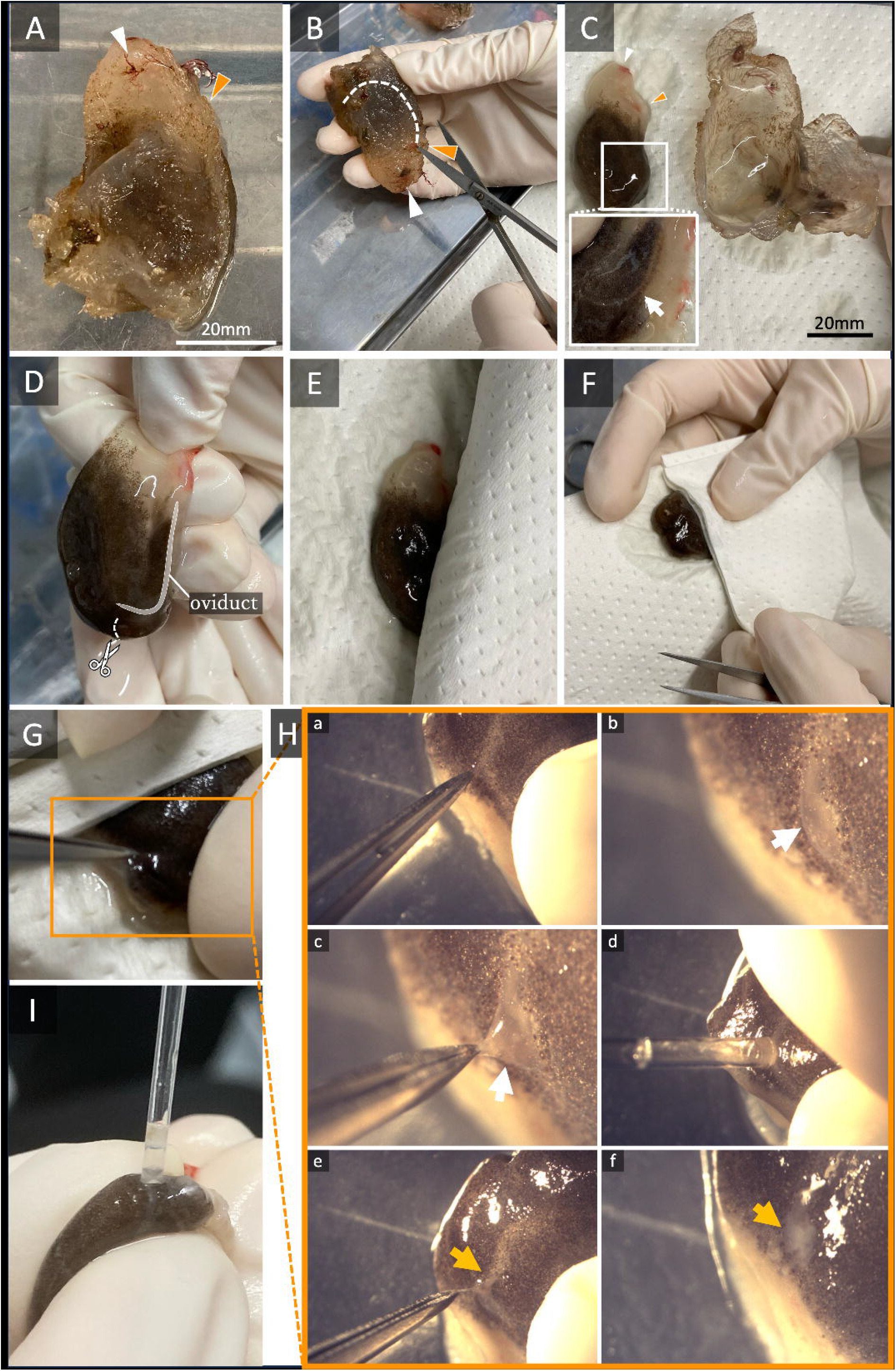
Dissection and isolations of eggs and sperm in *A. aspersa*. (A-B) *A. aspersa* adult. The white dotted line in B indicates the site of incision. White arrowhead: oral siphon. (C) *A. aspersa* without tunic. The white rectangle has been enlarged; the arrow indicates gonoduct. (D-G) Operational steps before collecting eggs and sperm. (Ha-f) Sequential photos during operation for eggs (Ha-d) and sperm (He-f) collection. Orange arrowhead: atrial siphon. Eggs in the oviduct can be observed (white arrow of Fig.1 Hc). The orange arrow indicates sperm duct and sperm. (I) Collecting of sperm with Pasteur pipette.

### 2.2 Isolation of eggs and sperm

Eggs and sperm isolations were performed following steps.

1. **Egg isolation:** Using micro-tweezers, carefully pick and tear the epithelial layer just above the oviduct (Fig. 1G) to expose the oviduct (Fig. 1Ha, Hb). Pinch off the oviduct without damaging the sperm duct, and create a small hole in the oviduct (Fig. 1Hc). Extract the eggs using a Pasteur pipette (Fig. Hd). Keep the specimen between paper towels (Fig. 1F-H) to prevent damage from acidic body fluid by avoiding sperm contamination.
2. **Sperm isolation:** Similarly, create a hole in the sperm duct using tweezers (Fig. 1He, Hf). Extract the sperm using a new Pasteur pipette (Fig. 1I) and transfer it to a new tube. Sperm can be stored for at least 24 hours at 4ºC.

### 2.3 Dechorionation of eggs

To observe embryonic development, the process of removing chorions and test cells from eggs, known as dechorionation, is essential.

1. **Dechorionation:** Immediately after isolating the eggs, the chorion and test cells surrounding the eggs were removed by the dechorionation solution, described in *Ciona* protocol (Hotta et al., 1999). In brief, the solution consisted of 0.05% actinase-E and 1% mercaptoacetic acid sodium salt in Millipore-filtered seawater (MFSW), stored at 4 °C. Before dechorionation, six drops of 2N NaOH were using a Pasteur pipette to 10 mL of the prepared solution in a glass tube and mixed well, followed by the addition of the collected eggs.
2. **Wash:** When 10% of the eggs have been dechorionated, thoroughly wash them with MFSW.
Note: If the dechorionation fails, change the amount of NaOH to adjust the pH. We don’t recommend dechorionation using the trypsin solution used for *Phallusia* (Sardet et al., 2011); in this case, transfer the oocytes individually to seawater immediately after dechorionation.
3. **Store:** Eggs can be stored in MFSW on 0.1% gelatin-coated dishes until fertilization (Funakoshi et al., 2021).

### 2.4 Fertilization

A high amount of sperm was used for fertilization as described below. This increased sperm concentration enables cleavage timing to be highly synchronized.

1. **Insemination:** Put dechorionated eggs in a 35 mm dish filled with 5 mL MFSW (Fig. 2A). Add 20 μL of undiluted dry sperm and mix well with MFSW.
2. Wait until approximately half of the eggs exhibit deformation due to sperm entry (Fig. 1B), typically observed within 5–10 min. The polar body can be observed after deformation (Fig. 1C, Roegiers et al., 1999)
3. **Wash:** Wash the eggs twice with MFSW and transfer them to a new 35 mL dish.
4. **Incubation:** Next, incubate the embryos according to the embryonic stages of *A. aspersa* provided in Funakoshi et al. (2020) and track the development of the embryos until the stage you intend to observe. Developmental speed until hatching is comparable to that of *Ciona*. For more details, you can refer to the RAMNe database here.

**Fig. 2.**
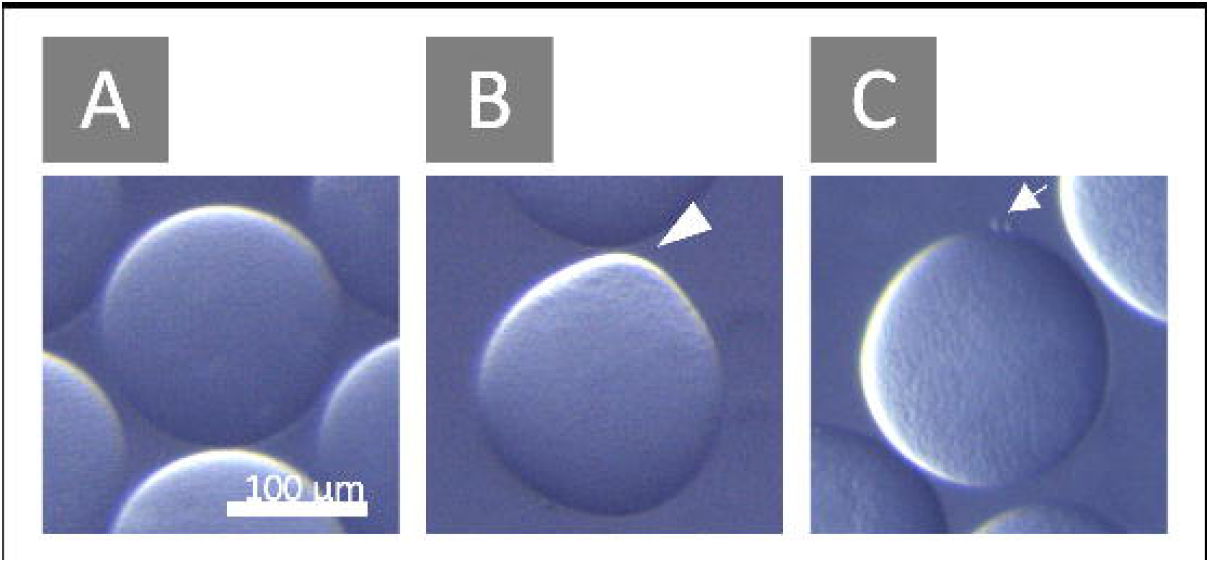
Example of egg deformation during fertilization. (A) Unfertilized egg. (B) Fertilized egg showing deformation. The white arrowhead indicates the contraction pole (Roegiers et al., 1999) (C) Meiosis resumes, and polar bodies become visible. The white arrow indicates the polar bodies.

### 2.4 Comparison of fertilization rate of Ascidiella aspersa with conventional protocols

1. **Fertilization rate:** To demonstrate the validity of this protocol, we compared the fertilization rate with the conventional fertilization protocol for *Ciona* and *Phallusia* (Christian Sardet et al. 2011). In particular, since *Phallusia* and *Ascidiella aspersa* belong to the same family, Ascidiidae (Shito et al., 2020), the series of protocols (dechorionation and fertilization) for *Phallusia* are usually considered applicable to *A. aspersa*. However, these protocols indicate only up to a 2% fertilization rate in *A. aspersa* (Table. 1, Fig. 3). On the other hand, the new protocol shows an 80% fertilization rate (Table. 1, Fig. 3).
2. **Incubation time:** Although the seawater appeared cloudy due to the large amount of sperm, no polyspermy occurred. In addition, we also confirmed that our new protocol is applicable to *Ciona* with a high efficiency of fertilization rate (data not shown), indicating that our protocol is generally applicable for other ascidian species. Our protocol also does not require preincubation times for activation or maturation of sperm (Fig. 4). Usually, preincubation of spermwas needed for efficient fertilization of ascidian eggs (C. Sardet et al. 1989). Our protocol demonstrates that *A. aspersa* eggs do not need such incubation time (Fig. 4). Typically, an incubation step in seawater is required after egg extraction and dechorionation, but even without this step, more than 60% of the eggs were successfully fertilized when inseminated immediately (Table 1).
3. **Validation of the Synchronous Fertilization:** Synchronous fertilization is possible with our new protocol. Fortunately, this beautiful transparent animal has the ability for mRNA translation before fertilization (Funakoshi et al. 2021). Thus, combined with high-rate fertilization protocol, it demonstrates the ability to observe some eggs’ dynamical signaling changes immediately after fertilization at the same time (Suppl. Mov. 1).

**Table 1.** Comparison of fertilization rate among different protocols. Form left column, conventional fertilization protocols of *Phallusia, Ciona* and New protocol in this study.

|  | *Phallusia*<br>protocol | *Ciona* protocol | New protocol |
| --- | --- | --- | --- |
| batch1 | - | - | 65 % |
| batch2 | 2.1 % | 1.8 % | 57 % |
| batch3 | 0 % | 0 % | 48 % |
| batch4 | 0.48 % | 0.92 % | 80% |

**Fig. 3.**
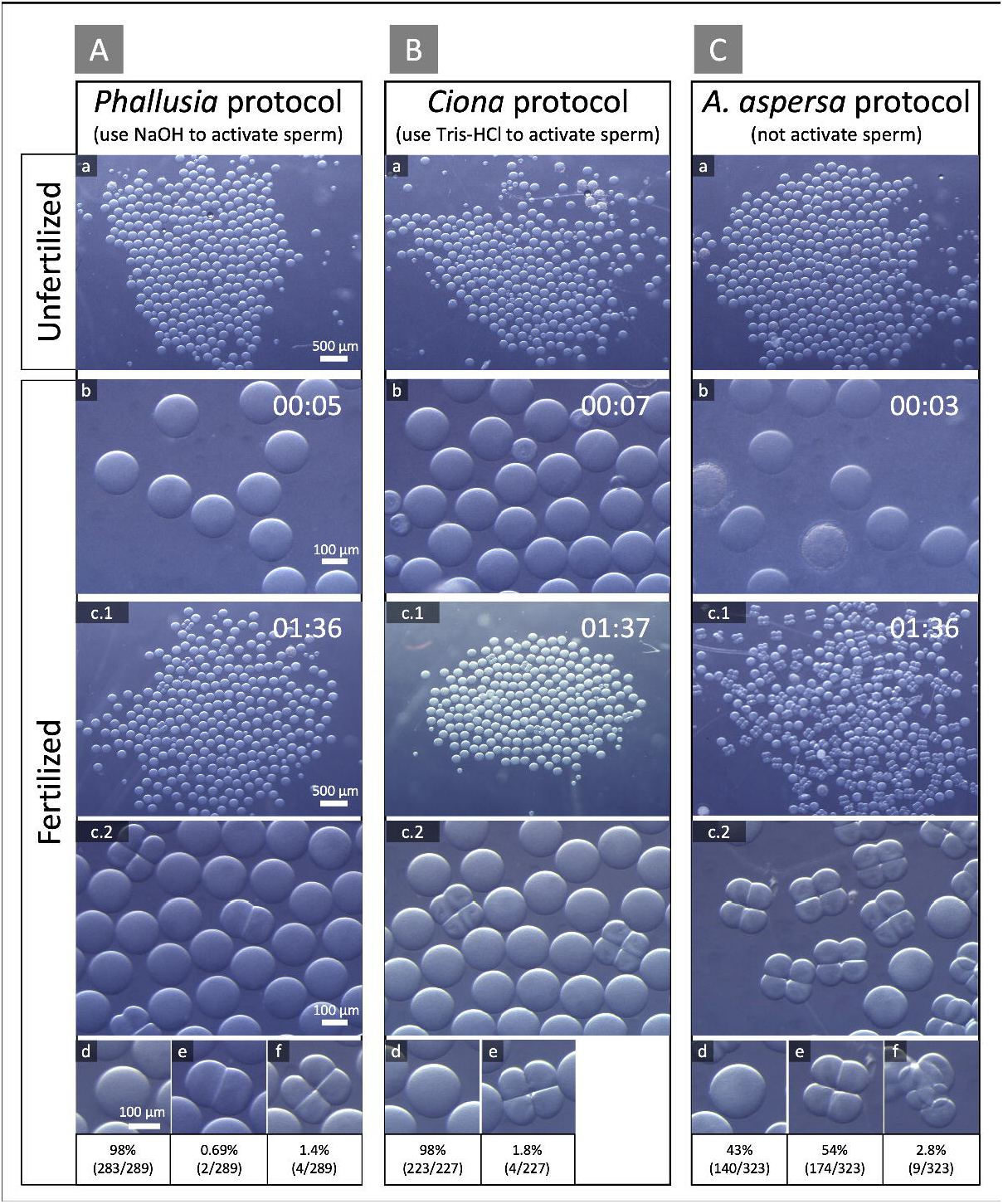
Comparison of the fertilization rates between conventional protocols (*Phallusia* and *Ciona*) and new *A. aspersa* protocol. The digits indicate the time after fertilization at 20 º C. The fertilization rates were about 2% and 60% in this batch for the conventional protocol and new protocol respectively.

**Fig. 4.**
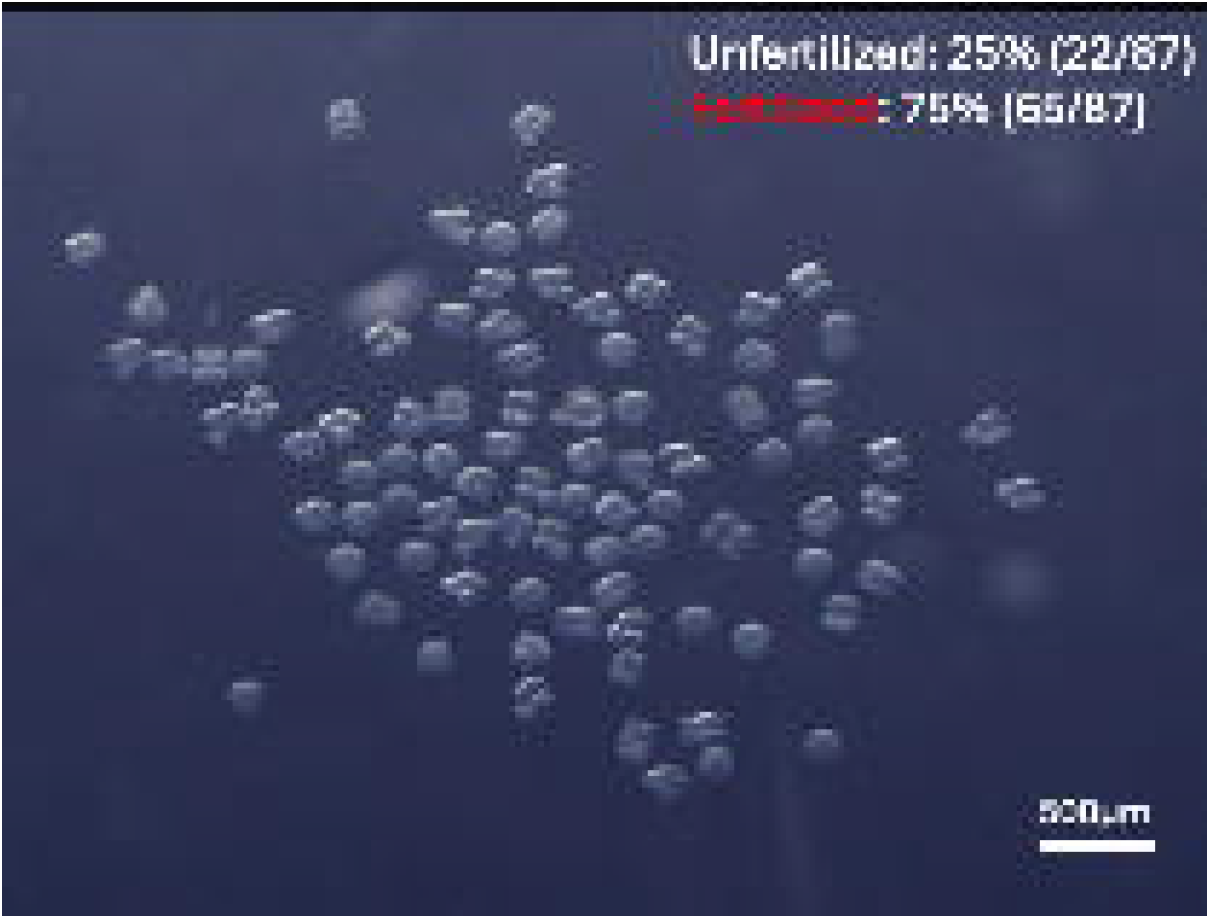
Fertilization rate and cleavage pattern when fertilized immediately after dechorionation.

## 3. Discussion

### 3.1 Advantages of new protocols

Our new protocol for fertilization consists of several steps, dissection, isolation, dechorionation, and fertilization. In each step, we have incorporated unique small improvements. In the dissection step, we pay attention to the contamination of the acidic body fluid with a paper towel (Fig. 1). In the isolation step, we adopted a method of directly collecting gametes from the gonoduct using a Pasteur pipette (Fig. 1Ha-f). In the dechorionation step, instead of trypsin, which is usually used in the same genus of *Ascidiella*, an actinase (pronase) solution used in *Ciona* and *Halocynthia roretzi* was applied. In the fertilization step, an unusually large excess of sperm was used for fertilization without activation, and the eggs were not pre-incubated in seawater.

These small improvements not only led to the success of improving the fertility rate of *A. aspersa*, a long-standing problem, but also brought the following benefits: high fertilization rate, high synchronization, a simple experimental procedure, short insemination time, and low polyspermy.

Especially, conventional fertilization protocols require dilution of sperm and subsequent activation by making the sperm alkaline to avoid polyspermy, which is a condition where multiple sperm penetrates an egg (Honegger and Koyanagi 2008). However, we have found that neither dilution nor activation of sperm is necessary to avoid polyspermy or to improve fertilization rates from below 10% to 80% (Table 1). The addition of this excess amount of sperm allows the eggs to be fertilized at a higher rate and enables highly synchronized cleavage timing (Fig. 4).

The fluctuation in egg fertilization rates can be due to condition of the eggs, such as being immature (see smaller oocytes) or damaged (opaqued). To increase the fertilization rate, the dechorionation protocol is also important to decrease oocyte damage. Long-time incubation in trypsin solution for dechorionation decreases the fertilization rate and might inhibit fertilization (Fuke and Numakunai 1999; Sawada and Saito 2022).

### 3.2 *A. aspersa*, to become a new model for chordate developmental studies

*A. aspersa* is distributed worldwide. Improved protocols will promote the use of this animal in experiments, making *A. aspersa* a potentially useful model organism. Developmental stages and morphological descriptions have already been established (Funakoshi et al. 2021). Whole genome sequencing and transcriptome analyses have also been performed by several groups. However, obtaining eggs remains a problem. Mature gonads are most abundant in winter and spring and least abundant in summer (Lynch et al. 2016). Optimizing laboratory rearing systems for this ascidian, such as those recently optimized for *Ciona*(Mathiesen et al. 2024), will further improve its utility as a model organism.

## 4. Materials and Methods

### 4.1 Sample preparation

*Ascidiella aspersa* is collected Onagawa field center or the Hakodate Fisheries Research Institute in Japan. *Animals are kept under 16 degrees in the tank until use*.

## Supporting information

Suppl. Mov. 1

## 5. Acknowledgements

Takumi T. Shito for handling of *A. aspersa* and valuable discussion.

Dr. Hitoyoshi Yasuo for checking protocol validity and valuable discussion.

Dr. Minoru Ikeda and Captain Toyokazu Hiratsuka (Onagawa Field Center of Tohoku University), Dr. Makoto Kanamori, Dr. Masafumi Natsuike, and Takuya Mizukami (Hokkaido Research Organization, Hakodate Fisheries Research Institute), Dr. Gaku Kumano (Graduate School of Life Sciences, Tohoku University), Mr. Akio Takiya, Dr. Tatsunari Mori, Dr. Takaaki Kayaba, and Dr. Motohito Yamaguchi (Hokkaido Research Organization, Central Fisheries Research Institute) for their help in collecting the samples.

## 8. Data Availability

The datasets presented in this study can be found in online repositories. The names of the repository/repositories and accession number(s) can be found in the article/Supplementary Material.

## 9. Figure legends

**Suppl. Mov. 1 Ca**^**+2**^ **wave at fertilization**

GCaMP6s mRNA was injected into unfertilized eggs to visualize Ca^2+^ elevation during fertilization. Our protocol enabled real-time imaging of synchronized fertilization events across multiple eggs.

